# Proteotyping SARS-CoV-2 virus from nasopharyngeal swabs: a proof-of-concept focused on a 3 min mass spectrometry window

**DOI:** 10.1101/2020.06.19.161000

**Authors:** Duarte Gouveia, Guylaine Miotello, Fabrice Gallais, Jean-Charles Gaillard, Stéphanie Debroas, Laurent Bellanger, Jean-Philippe Lavigne, Albert Sotto, Lucia Grenga, Olivier Pible, Jean Armengaud

## Abstract

Rapid but yet sensitive, specific and high-throughput detection of the severe acute respiratory syndrome coronavirus 2 (SARS-CoV-2) in clinical samples is key to diagnose infected people and to better control the spread of the virus. Alternative methodologies to PCR and immunodiagnostic that would not require specific reagents are worth to investigate not only for fighting the COVID-19 pandemic, but also to detect other emergent pathogenic threats. Here, we propose the use of tandem mass spectrometry to detect SARS-CoV-2 marker peptides in nasopharyngeal swabs. We documented that the signal from the microbiota present in such samples is low and can be overlooked when interpreting shotgun proteomic data acquired on a restricted window of the peptidome landscape. Simili nasopharyngeal swabs spiked with different quantities of purified SARS-CoV-2 viral material were used to develop a nanoLC-MS/MS acquisition method, which was then successfully applied on COVID-19 clinical samples. We argue that peptides ADETQALPQR and GFYAQGSR from the nucleocapsid protein are of utmost interest as their signal is intense and their elution can be obtained within a 3 min window in the tested conditions. These results pave the way for the development of time-efficient viral diagnostic tests based on mass spectrometry.

## Introduction

The new severe acute respiratory syndrome-related coronavirus 2 (SARS-CoV-2) is the causative agent of COVID-19, the coronavirus disease that was first reported in December 2019 in the city of Wuhan, China (1). Due to its easy inter-human transmission, SARS-CoV-2 has since quickly spread worldwide, causing more than 8 million COVID-19 diagnosed infections and more than 450 thousand deaths officially reported as off mid June 2020 (https://covid19.who.int/). The rapid, sensitive and specific detection of the SARS-CoV-2 virus in large cohorts of clinical samples is of utmost importance to identify infected people and control the propagation of the virus by specific containment measures. At the same time, being able to catch the numerous SARS-CoV-2 variants represents an opportunity to identify attenuated forms of the virus (2). However, the occurrence of specific mutations, especially deletions, may challenge current molecular detection methodologies.

The research community has been placing great efforts in the development of quick and accurate detection tests (3, 4). The gold standard in diagnostics relies on the amplification and measurement of the viral RNA by reverse transcription polymerase chain reaction (RT-PCR). RT-PCR is highly specific and achieves a good compromise between speed (90-300 min) and sensitivity. However, due to the great demand for PCR-based testing, shortage of RNA extraction kits and PCR reagents may have limited the testing capacity in some countries at the early stage of the pandemic (5). Besides, RT-PCR testing of clinical samples may be in some case less efficient due to nucleic acid variations in the targeted regions - primers or their close vicinity - that could affect the amplification rate (6, 7). For these reasons, alternative detection strategies that address these concerns should be developed to complement conventional tools.

Immunoassays, whole-genome sequencing (8) and mass spectrometry (MS) (9) technologies are commonly suggested alternatives to PCR-based assays. Among these, new generation MS offers a highly sensitive technology that allows the rapid identification of thousands of proteins present in a single sample. The typing of organisms by tandem mass spectrometry (MS/MS), commonly referred to as “proteotyping”, is based on the identification of specific peptide sequences that allow the unambiguous identification of organisms (9–11). The uniqueness of the mass to charge ratios and fragmentation patterns measured in MS/MS allows identifying peptides that differentiate organisms at the subspecies level. Although classical MS-based identification of pathogens in the clinical setting is based on whole-cell MALDI-TOF technology (12), the field has thrived with the increases in speed, sensitivity, and accuracy of new MS instrumentation in the last decade. The coupling of new generation instruments with the separation power of liquid chromatography makes LC-MS/MS a valuable technology to implement in the routine of clinical laboratories. Despite their high potential, the application of LC-MS/MS approaches for virus proteotyping is still scarce. Among the few examples available in the literature, LC-MS/MS was shown to be able to detect purified influenza virus (13) and human metapneumovirus in clinical samples (14).

Because of the considerable damages of the COVID19 pandemic, the mass spectrometry community quickly proposed to mobilize its efforts at helping to understand the molecular mechanisms of infection (15–17) and at improving detection methods (18). Several research groups started investigating MS-based quantification of peptides for the detection of SARS-CoV-2 in clinical samples, but these results are not yet published (19–22). These preliminary results indicate that targeted MS, in which the mass spectrometer is programmed to precisely detect and quantify a limited number of peptides of interest, can be successfully applied to virus detection. Targeted MS is considered as the gold standard for peptide quantification due to its higher sensitivity when compared to shotgun proteomics approaches. Nevertheless, this approach has a much lower throughput and is commonly used to test hypotheses on a subset of proteins of interest, in contrast to discovery shotgun proteomics. By being more flexible, the latter provides a more comprehensive picture of the viral peptidome, including the detection of variant sequences because of the possibility of detecting peptides without any previous knowledge of their sequences.

Here we established the proof of concept of the use of MS/MS for the rapid proteotyping of SARS-CoV-2 from clinical samples. We recently published a dataset from a shotgun LC-MS/MS experiment performed with SARS-CoV-2 infected cells and proposed a list of specific viral peptides that could be used for the development of targeted approaches (23). Interestingly, we observed that some SARS-CoV-2 specific peptides eluted from the LC column at narrow windows of retention time. Here, we used LC-MS/MS with an Orbitrap instrument (Q Exactive HF) for analyzing the peptidome from nasal swabs spiked with different quantities of viral material. By using a short LC gradient focusing on the region of interest identified in our previous study, we tested the detection of the virus in samples containing different quantities of viral peptides, as well as COVID-19 clinical samples, paving the way for the development of time-efficient viral diagnostic tests based on an alternative platform.

## Materials and Methods

### Nasopharyngeal swab collection and processing for reference matrices

Two nasopharyngeal swabs were collected using a sterile polyester swab with semi-flexible polystyrene handle (Puritan) from two healthy volunteers (swabs R1 and R2). Each swab was soaked into a tube containing 200 μL of sterile water, incubated for 10 min at room temperature, and then rinsed with 200 μL of sterile water. The biological material from the 400 μL of solution was precipitated with the addition of 100 μL of trichloroacetic acid at 50% (w/v) and centrifugation at 16,000 g for 10 min. The supernatant was discarded. The hardly visible pellet was dissolved into 50 μL of LDS1X containing 5% beta-mercaptoethanol, heated for 5 min at 99°C, and centrifuged briefly. For each swab sample, a volume of 20 μL of LDS1X sample was deposited on a SDS-PAGE gel and run for 5 min. After migration, the gel was rinsed with water, stained with Simply Blue SafeStain (Invitrogen), and destained overnight in water. The two polyacrylamide gel bands corresponding to the whole proteome of each matrix were excised, processed as described (24), and then subjected to trypsin Gold proteolysis (Promega) using 0.01% ProteaseMAX surfactant (Promega). The nasal matrix peptide fractions were 50 μL for each swab.

### NanoLC-MS/MS characterization of the peptides extracted from the nasopharyngeal swab matrices

Peptides from the nasal swab matrices were analysed with a Q-Exactive HF mass spectrometer (Thermo) coupled with an UltiMate 3000 LC system (Dionex-LC) and operated in data-dependent mode as previously described (25). A volume of 4 μL of peptides was injected, desalted onto an Acclaim PepMap100 C18 pre-column (5 μm, 100 Å, 300 μm id × 5 mm), and then resolved onto a nanoscale Acclaim PepMap 100 C18 column (3 μm, 100 Å, 75 μm id × 50 cm) with a 90-min gradient at a flow rate of 0.2 μL/min. The gradient was developed from 4 to 25% of CH3CN, 0.1% formic acid in 75 min, and then from 25 to 40% in 15 min, washed, and re-equilibrated. Peptides were analysed with scan cycles initiated by a full scan of peptide ions in the Orbitrap analyser, followed by high-energy collisional dissociation and MS/MS scans on the 20 most abundant precursor ions (Top20 method). Full scan mass spectra were acquired from *m/z* 350 to 1500 at a resolution of 60,000 with internal calibration activated on the *m/z* 445.12002 signal. Ion selection for MS/MS fragmentation and measurement was performed applying a dynamic exclusion window of 10 sec and an intensity threshold of 5×10^4^. Only ions with positive charges 2+ and 3+ were considered.

### Cell culture and Virus

Vero E6 (ATCC, CLR-1586) cells were cultured at 37°C in 9% CO_2_ in Dulbecco's modified Eagle's medium (DMEM, Gibco, ThemoFisher) supplemented with 5% fetal calf serum (FCS) and 0.5% penicillin–streptomycin. The SARS-CoV-2 strains 2019-nCoV/Italy-INMI1 (Genbank MT066156) was provided by the Lazzaro Spallanzani National Institute of Infectious Diseases (Rome, Italy) via the EVAg network (European Virus Archive goes global). SARS-CoV-2 stocks used in the experiments had undergone two passages on Vero E6 cells and were stored at - 80°C. Virus titer was 7.25×10^5^ plaque forming units (PFU)/mL, as determined by standard plaque assay (three dilutions in duplicates). All experiments entailing live SARS-CoV-2 were performed in our biosafety level 3 facility and strictly followed its approved standard operating procedures.

### Infection and virus purification

Vero E6 cells (1×10^6^) seeded into 150 cm^2^ flasks were grown to cell confluence in 15 ml DMEM supplemented with 5% FCS and 0.5% penicillin–streptomycin for one night at 37°C under 9% CO_2_. They were infected at multiplicity of infection (MOI) of 0.001. Cells were harvested at 3 days post infection (dpi) and viral suspension was recovered after centrifugation at 2,500 rpm for 5 min to remove cell debris. 33 ml of the viral suspension were laid on 5 ml of 20% (w/v) sucrose cushion prepared in NaCl 0.1 M, EDTA 1m M, 10 mM Tris HCl buffer (pH 7.4) (TNE buffer) in Ultra-Clear 38 ml tubes (Beckman Coulter). Samples were centrifuged at 25,000 rpm for 2h at 4°C. Pellets were solubilised in 150 μL of cold TNE buffer and a volume of 1.5 ml was laid on a five step 20 - 60% (w/v) sucrose gradient prepared in Ultra-Clear 13 ml tubes (Beckman Coulter). The tubes were centrifuged at 35,000 rpm for 2h at 4°C in a Beckman SW41 rotor. After recovery of the virus band, the viral suspension was inactivated by incubation with betapropiolactone at a final concentration of 0.5% for 72h at 4°C. Plaque assay titration was used to quantify the purified virus and validate the viral inactivation.

### Proteolysis of purified SARS-CoV-2 virus

The inactivated purified virus sample (equivalent to 7.25×10^5^ PFU/mL) was quantified in terms of protein concentration (0.614 mg/mL) by UV spectrophotometry. A volume of 60 μL was mixed with 20 μL of LDS3X to obtain a protein fraction of 0.46 mg/mL. After denaturation at 99°C for 5 min, a volume of 25 μL (11.5 μg of proteins) was deposited on a NuPAGE 4-12% gel (Invitrogen) and subjected to 5 min electrophoretic migration. The whole proteome was excised as a single polyacrylamide gel band and subjected to trypsin proteolysis as previously described (24). An aliquot of 50 μL of peptides was extracted. MS/MS analysis was performed to confirm the high content of viral proteins in this sample (data not shown).

### Preparation of simili SARS-CoV-2 contaminated swabs

SARS-CoV-2 viral peptides (3 μL) were diluted in 6 μL of H_2_O, 0.1% TFA. After mixing, 3 μL of this tube was removed and diluted with 6 μL of H2O, 0.1% TFA. This was repeated several times to obtain a one third dilution cascade of viral peptides. Two series of simili swabs were prepared in parallel. The two peptide fractions obtained from nasopharyngeal swabs (35μL) were diluted with 15 μL of H_2_0, 0.1% TFA. A volume of 6 μL of this diluted matrix was added to each simili swab samples, giving a final volume of 12 μL per sample. Thus, each simili swab contained the equivalent of 8.4% of the proteins harvested by a nasal swab. Two biological replicates were prepared using each nasal swab matrix. A volume of 10 μL per sample was injected in the Q-Exactive HF tandem mass spectrometer. They were analysed in the same conditions as above except that the gradient was developed from 8 to 12.5% of CH3CN, 0.1% formic acid for 30 min at a flow rate of 0.2 μL/min. The 20 min MS/MS acquisition started 17 min after injection.

### COVID-19 nasopharyngeal swabs and MS/MS measurements

Nasopharyngeal swabs were collected from COVID-19 diagnosed adult patients as routine medical controls and tested by RT-PCR assay for detecting SARS-CoV-2 in a nasopharyngeal sample (swabs T1-T9). This study was approved by the institutional review boards of the University Hospital of Nîmes, France (2020-05-01). Patients have been previously be informed that part of these samples could be used for research purpose and agreed. Each swab was soaked into a tube containing 5 mL of Phosphate Buffered Saline (pH 7.4) sterile solution and transferred in the biosafety level 3 facility. The biological material was precipitated with the addition of 1.25 mL of trichloroacetic acid at 50% (w/v). After centrifugation, the supernatant was discarded and the pellet was dissolved into 25 μL of LDS1X containing 5% beta-mercaptoethanol, heated for 5 min at 99°C, and deposited on a NuPAGE 4-12% gel (Invitrogen). The proteins were subjected to 5 min electrophoresis and treated as described here above to obtain tryptic peptides. MS/MS acquisition was done as for the simili swabs. The 20 min MS/MS acquisition started 17 min after injection with an inclusion list comprising 28 *m/z* values corresponding to 23 viral peptides.

### Peptide assignation and proteomics data analysis

MS/MS spectra from the nasopharyngeal swabs were searched against the generalist NCBInr database (108,307,546 sequences totalling 41,817,980,956 amino acids) with the MASCOT Daemon 2.3.2 search engine (Matrix Science). The search parameters were as follows: full-trypsin specificity, maximum of two missed cleavages, mass tolerances of 5 ppm on the parent ion and 0.02 Da on the MS/MS, carbamidomethylated cysteine (+57.0215) as a fixed modification, and oxidized methionine (+15.9949) and deamidation of asparagine and glutamine (+0.9848) as variable modifications. PSMs with an FDR<1% were selected for peptide inference. Peptides were assigned to taxa using the Unipept 4.3 web interface (26) with default parameters (equate I/L, filter duplicate peptides).

MS/MS spectra from the simili SARS-CoV-2 contaminated swabs and from the COVID-19 nasopharyngeal swabs were assigned with the MASCOT Daemon 2.3.2 search engine (Matrix Science) as follows: the spectra were first queried against the cRAP_contaminants_2020-05-18.fasta file and then against the Swissprot_Human_ISL_410545_2020-05-18 database (20,139 sequences totalling 11,330,214 amino acids) in follow-up mode and with the decoy option activated. This last database is the merge of the SARS-CoV-2 viral proteins and the Swissprot Human proteome. The MASCOT search was performed with the same parameters as above. All peptide matches presenting a MASCOT peptide score with a FDR lower than 1% were assigned to protein sequences. MS1 peak areas were evaluated with Skyline (27). Briefly, we created spectral libraries based on the DAT files from each MASCOT search (cut-of 0.99) and uploaded the MS1 full scan information contained in the raw files. The protein database previously used for the MASCOT search was used as background proteome. Only the viral proteins were added to the target panel. Peptide settings were matched to those used in the MASCOT search. Peak peaking was manually checked for all peptides.

### Mass spectrometry and proteomics data

The mass spectrometry and proteomics data acquired on simili swabs have been deposited to the ProteomeXchange Consortium via the PRIDE partner repository (28) with the dataset identifiers PXD019686 and 10.6019/PXD019686.

## RESULTS

### A shotgun MS-based strategy elaborated with simili COVID-19 swabs

To assess the performance of shotgun MS-based proteomics in detecting SARS-CoV-2 peptides in a background matrix consisting of nasopharyngeal swab protein material, we experimentally created tryptic peptidomes from i) a purified virus solution obtained from Vero E6 cells infected with a SARS-CoV-2 reference strain, and ii) nasopharyngeal swabs obtained from two healthy volunteers **(Figure 1)**. We first characterized the nasal peptidomes and searched for the presence of detectable microorganisms by metaproteomic data analysis. Then, the virus peptidome was serially diluted into nasopharyngeal swab peptidomes to obtain two sets of seven tubes containing from 460 ng (equivalent to 544 infectious particles) to 0.6 ng (equivalent to 1 infectious particle) of viral protein material. The fourteen samples were subsequently analyzed by LC-MS/MS. A window of 20 min of acquisition within a 30 min LC gradient was adjusted to target the region of elution of five previously identified virus-specific peptides (23). The rationale for focussing the mass spectrometry measurements on these peptides was their remarkable sequence conservation amongst the numerous SARS-CoV-2 strains sequenced to date or/and their specificity to the novel coronavirus (23). These peptides were the following: EITVATSR, GFYAEGSR, HTPINLVR, IAGHHLGR, and ADETQALPQR. While known variants exist for the latter, the other four peptides are conserved along the several SARS-CoV-2 sequenced genomes.

**Figure 1.**
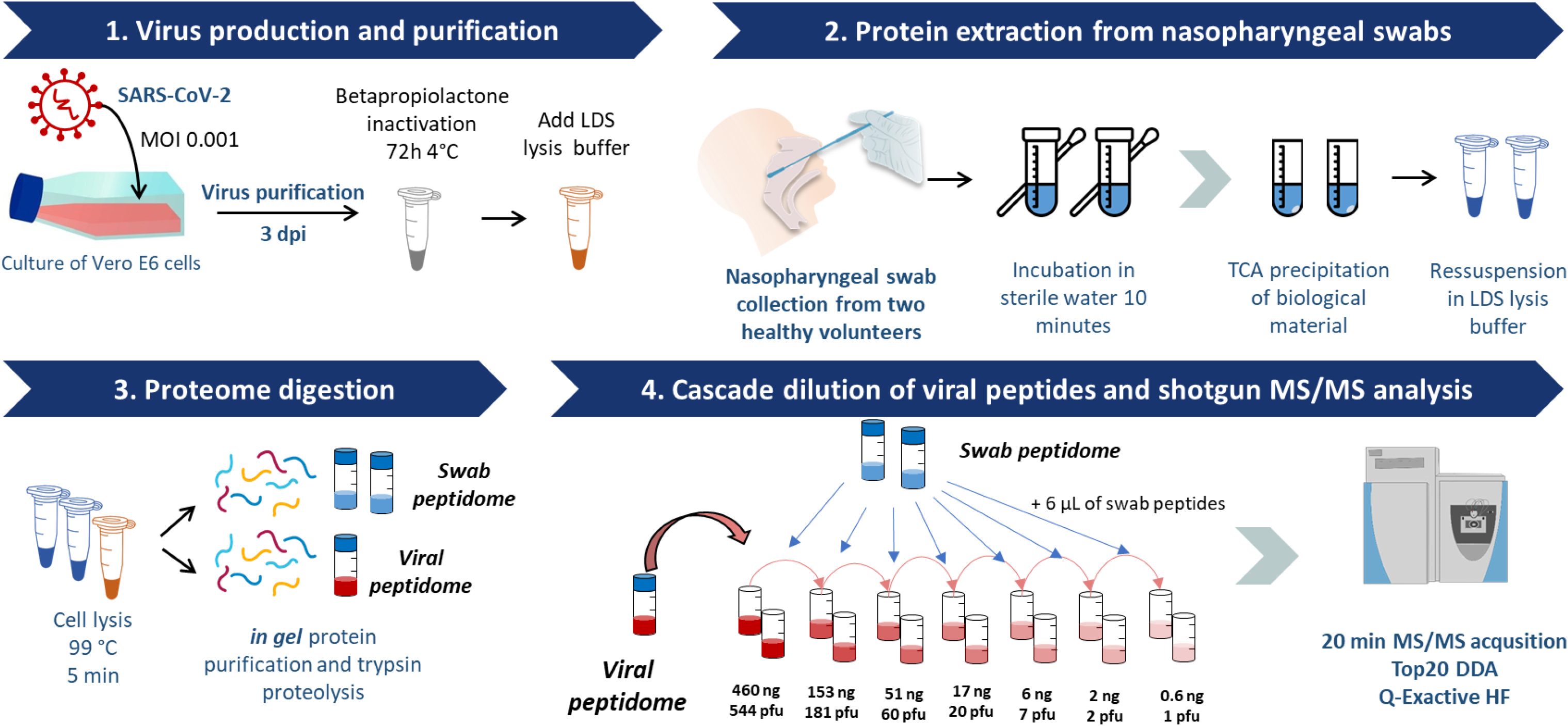
Strategy for the analysis of simili SARS-CoV-2 swabs (from nasal swab to MS/MS measurements).

### Metaproteomics analysis of the two nasopharyngeal swabs excludes the presence of abundant microorganisms

The MS/MS spectra acquired over 90 min on the two nasopharyngeal swabs were analyzed to infer the main microbial components present in these samples as such presence should be taken into account for creating an ad hoc database for MS/MS interpretation. Swabs R1 and R2 yielded 60,932 and 61,053 MS/MS spectra, respectively, from which 13,602 and 13,802 were attributed to 6,460 and 6,421 peptide sequences from organisms present in the NCBInr database (FDR <1%). These peptide sequences were analyzed with the Unipept tool (26) to assess the biodiversity present in each sample through their taxon-specificity characteristics based on the lowest common ancestor approach. Only a small proportion of the peptide sequences mapped by Unipept belonged to microorganisms (**Table S2)**. A rather low number (38 and 69) of peptides from the R1 and R2 swabs, respectively, were attributed to Bacteria, Archaea or Fungi. These corresponded to 0.6 % and 0.9% of the mapped peptide sequences, respectively. To exclude false positive identifications, we applied a threshold of at least three-taxon specific peptides for organism validation at the species level, corresponding to 0.5% of the total number of species-specific peptides (405 and 454 in each sample), as suggested by (29). Thus, one low-abundant *Corynebacterium* was identified in sample R1, namely *Corynebacterium accolens*, with 6 specific peptides. In swab R2, *Corynebacterium propinguum, Corynebacterium pseudodiphtheriticum,* and *Dolosigranulum pigrum* could be identified at the species taxonomical rank with 4, 5 and 15 specific peptide sequences, respectively.

### Detection of SARS-CoV-2 viral peptides in the simili swabs

Simili swabs containing specific quantities of SARS-CoV-2 virus and the equivalent of 8.4% of the nasal matrix protein material collected during sampling were analysed by MS/MS with a short gradient. We first confirmed on the most diluted fraction that the bacterial signal was negligible for both fractions, thus not to consider at the MS/MS attribution search stage. For this, the two datasets were searched against the generalist database NCBInr to check for the presence of non-human peptides in the swab peptidomes. The Unipept analysis of the detected peptide sequences showed that only 3 and 2 peptides, from replicate 1 and 2, respectively, were attributed to Bacteria and no bacterial species could be confidently identified (**Table S2**).

The results from the short gradient MS analysis on the simili swabs against the specific human/virus database yielded 139,404 MS/MS spectra recorded in the fourteen samples. From these, 34,647 were attributed to 2,919 peptide sequences with a FDR below 1% (**Table S3**). This data allowed for the identification of 1,094 protein groups (**Table S4**). A small fraction of 173 peptide-to-spectrum matches (PSMs), corresponding to 0.5% of the total PSMs, allowed identifying 18 different viral peptide sequences, including the five peptides of interest. The 18 peptides report for 3 structural proteins from the virus: 8 peptides from the nucleocapsid protein (N), 7 peptides from the spike protein (S), and 3 peptides from the membrane glycoprotein (M).

At least one viral peptide was identified in all samples independently of the concentration of the viral material, from 460 ng (544 PFU) to 2 ng (2 PFU). However, no peptide from the virus was identified in the sample containing viral peptides corresponding to 0.6 ng (1 PFU). The heatmap in **Figure 2** displays the MS1 peak areas, the number of PSMs attributed to each peptide in each sample, and the number of viral peptides identified in each sample. The five peptides with the highest MS1 peak areas across samples were the following: EITVATSR, GFYAEGSR, LNQLESK, ADETQALPQR, and KADETQALPQR. Among them, EITVATSR, GFYAEGSR, and ADETQALPQR are between the five peptides of interest. Peptide HTPINLVR was the seventh most abundant. Inversely, peptide IAGHHLGR was amongst the peptides with the lowest MS1 peak areas, along with peptides MSECVLGQSK, LDDKDPNFK, and EIDRLNEVAK.

**Figure 2.**
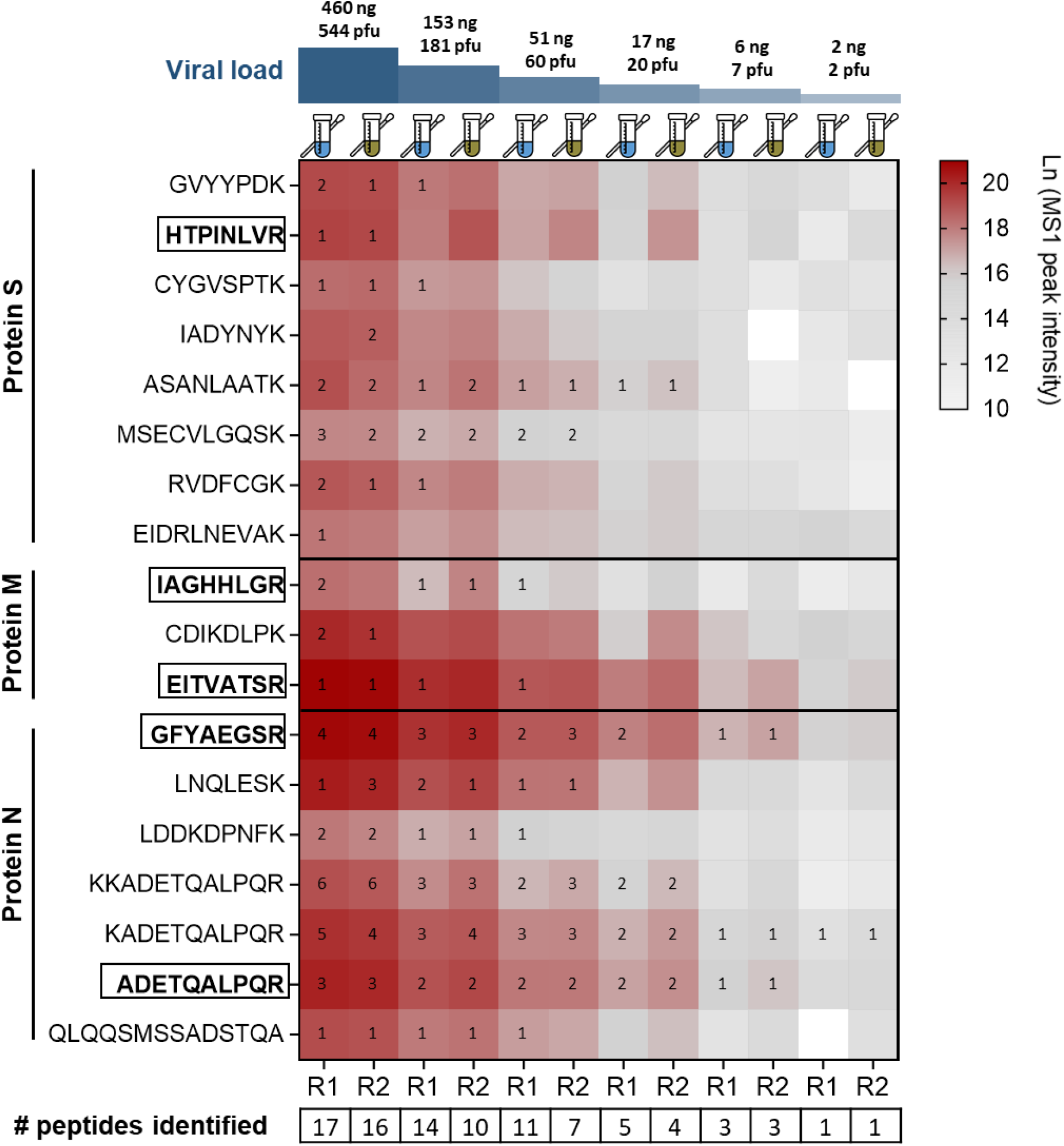
Heatmap of peptide intensities in the samples containing different viral loads. Cell colour corresponds to MS1 peak area, red being the highest and white the lowest. Numbered cells correspond to the number of PSMs from the MS/MS search that identified the peptide; cells with zero values mean that no MS/MS spectra was attributed to the peptide in that sample (at FDR 1%). The four peptides of interest are in bold and squared. The number of identified peptides in each sample is indicated on the bottom of the figure. R1 and R2 stand for “replicate 1” done with nasopharyngeal matrix 1 and “replicate 2” done with matrix 2 from each viral load condition. Viral load is given by the quantity of viral protein material contained in each sample (in ng), and the number of estimated infectious viral particles (in PFU).

As expected, the number of identified peptides decreased with the decreasing viral load in the sample. While all 18 peptides were identified in the initial dilution containing 460 ng of viral proteins (544 PFU), in highly-diluted samples containing 6 ng of viral proteins (7 PFU) and 2 ng (2 PFU), the virus was proteotyped with only 3 and 1 peptides, respectively. In these samples only peptides from protein N were detected (ADETQALPQR, KADETQALPQR, and GFYAEGSR). Generally, the peptides from protein N were the most consistently detected across samples. Despite being among the peptides with higher peak areas in the chromatograms, peptides of interest HTPINLVR and EITVATSR were only detected in the simili swabs containing an estimated 460 ng of viral proteins (544 PFU). On the other hand, the two other peptides of interest from protein N, GFYAEGSR and ADETQALPQR, allowed virus proteotyping in the sample containing 6 ng of viral proteins (7 PFU). Of note, the peptide identified in the condition with 2 ng of viral proteins (2 PFU) is a miss-cleaved version of the ADETQALPQR peptide: KADETQALPQR. Peptide ASANLAATK, that had not been previously selected among the “best” candidates for SARS-CoV-2 proteotyping (23), was detected in the dilution with 17 ng of viral proteins (20 PFU) and was the most sensitive peptide from protein S.

**Figure 3** represents the retention times of viral peptides from 2 to 19 min of MS acquisition, and their intensities in the samples with the highest concentration of viral proteins (460 ng, 544 PFU). Peptides of interest are squared and have well visible peaks. With the LC gradient used in this experiment and the delay for starting the acquisition, the retention times of peptides are minus 4-6 minutes compared to those described in our previous paper. Peptides are generally well distributed along the gradient, with some exceptions of peptide pairs that co-elute: KADETQALPQR/RVDFCGK, QLQQSMSSADSTQA/CYGVSPTK, ADETQALPQR/GFYAQGSR, or HTPINLVR/EIDRLNEVAK. Six peptides elute in the first ten minutes of the gradient: IAGHHLGR, KKADETQALPQR, ASANLAATK, LNQLESK, KADETQALPQR, and RVDFCGK. Of the utmost interest, three of the most conserved and well-detected peptides, EITVATSR, ADETQALPQR, and GFYAQGSR, elute in a 3-min window between 13 and 16 min of the gradient.

**Figure 3.**
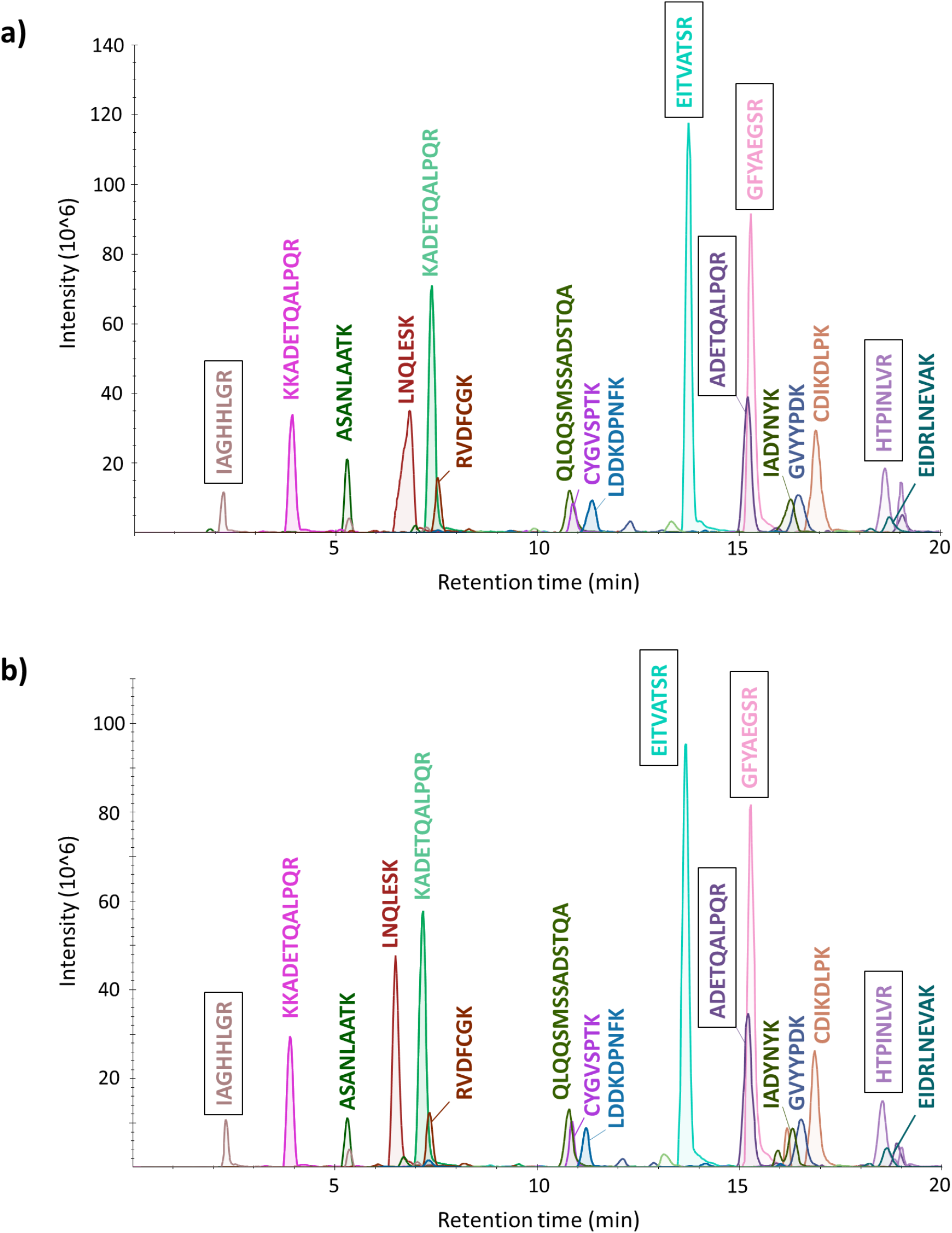
A view on the retention times of the viral peptides detected in the most concentrated simili swabs containing 460 ng of viral material (544 PFU). Nasopharyngeal matrix R1 (panel a) and matrix R2 (panel b).

### Detection of SARS-CoV-2 in clinical nasopharyngeal swab samples from COVID-19 diagnosed patients

Nasopharyngeal swabs were sampled from nine COVID-19 diagnosed patients with different clinical manifestations (moderate symptoms and asymptomatic) and at different post-diagnostic stages (**Table 1**). Due to the complexity of the samples, an inclusion list of *m/z* signals corresponding to the five peptides of interest as well as other SARS-CoV-2 peptides detectable in this gradient region (17) was added to the acquisition method to increase the likelihood of their detection. **Table S1** reports this inclusion list which contained *m/z* values for 28 different precursors from 23 different viral peptide sequences. The short gradient MS analysis on these clinical samples yielded between 655 and 1,151 MS/MS spectra recorded per sample. Sixty-five spectra were attributed to viral peptide sequences with a FDR below 1% (**Table S5**). This data allowed for the detection of six peptides reporting for two viral proteins (**Table S6**): LDDKDPNFK, KADETQAIPQR, KKADETQAIPQR, ADETQAIPQR, GFYAEGSR from protein N, and EITVATSR from protein M. The heatmap in **Figure 4** displays the MS1 peak areas, the number of PSMs attributed to each peptide in each sample, the number of viral peptides identified in each sample, and the result from the PCR testing performed on the same sample. The virus was confidently proteotyped in clinical swabs T7 and T8, with four and five peptides respectively. Peptide EITVATSR was identified in swab T4 with two spectral counts, but virus detection in this sample cannot be validated since this peptide is not specific to SARS-CoV-2 (23). As shown in **Table 1**, swabs T7 and T8 correspond to patients that were diagnosed as SARS-CoV-2 positive by RT-PCR with relatively clear viral loads (CT values of 28 and 25, respectively) and were sampled 11 and 4 days after their diagnostic and confinement. MS/MS samples were negative for swabs that yielded a relatively low PCR signal (CT of 35 and 36 for swabs T9 and T5), with undetectable PCR signal (swabs T2, T3, T4, and T6) or 14 days after their diagnostic (swab T1). The negative MS/MS signals for swabs T9, T5, and T1 patients are explained by the very low viral load probably present in these samples.

**Table 1.** COVID-19 nasopharyngeal swab medical samples.

| COVID-19 swab sample | Patient age | Pathology severity (*) | Days post-confinement | PCR control (#) | MS/MS |
| --- | --- | --- | --- | --- | --- |
| Swab T1 | 58 | Moderated | 14 | CT 23 | Negative |
| Swab T2 | 79 | Asymptomatic | 21 | Negative | Negative |
| Swab T3 | 68 | Moderated | 11 | Negative | Negative |
| Swab T4 | 91 | Asymptomatic | 10 | Negative | Negative |
| Swab T5 | 87 | Asymptomatic | 13 | CT 36 | Negative |
| Swab T6 | 75 | Asymptomatic | 11 | Negative | Negative |
| Swab T7 | 79 | Moderated | 11 | CT 28 | Positive |
| Swab T8 | 96 | Asymptomatic | 4 | CT 26 | Positive |
| Swab T9 | 94 | Moderated | 21 | CT 36 | Negative |

(\*) Moderated severity with radiological visible signs or asymptomatic
(#) PCR control done within 24h after control sampling

**Figure 4.**
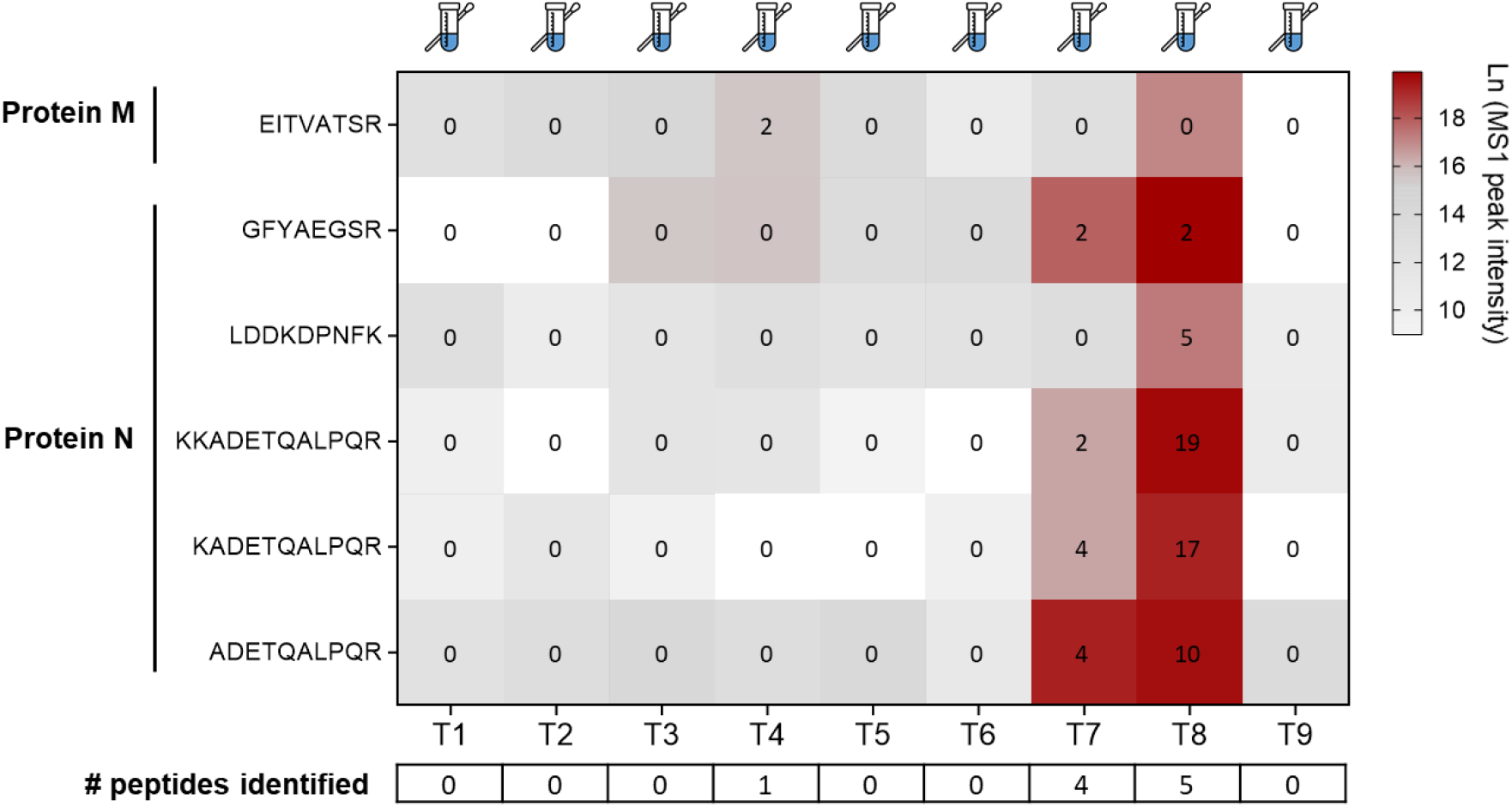
Heatmap of peptide intensities in the clinical nasopharyngeal swabs. Cell colour corresponds to MS1 peak area, red being the highest and white the lowest. Numbered cells correspond to the number of PSMs from the MS/MS search that identified the peptide; cells with zero values mean that no MS/MS spectra was attributed to the peptide in that sample (at FDR 1%). The number of identified peptides in each sample is indicated on the bottom of the figure. Patients were numbered from “swab T1” to “swab T9”.

## DISCUSSION

To foster the development of alternative detection methods for SARS-CoV-2, we performed a proof-of-concept study to assess the potential of MS/MS for proteotyping SARS-CoV-2: i) in simulated nasal swabs containing different quantities of viral peptides; and ii) in nasopharyngeal swabs from COVID-19 diagnosed patients. The two nasal peptidomes collected from healthy donors for the first experiment were first analyzed with a gradient of 90 min to check for the presence of detectable microorganisms from the natural microbiota. A search against a generalist database such as NCBInr detected only trace levels of very low abundant bacteria commonly found in the nasal tract (30), thus confirming the absence of a measurable microbiome in the swab samples. Based on this metaproteomic analysis, we used a human-only database as representative of the nasopharyngeal matrices for the subsequent analysis.

The simili SARS-CoV-2 contaminated swabs contained a fixed amount of swab peptidome, plus a precise amount of viral peptidome corresponding to the expected quantities extracted from 460 ng (544 PFU), 153 ng (181 PFU), 51 ng (60 PFU), 17 ng (20 PFU), 6 ng (7 PFU), 2 ng (2 PFU), and 0.6 ng (1 PFU) of SARS-CoV-2. It is important to note that the virus produced in Vero E6 cells and purified on sucrose gradient is only partially infectious, and thus the data are also presented in quantities of viral proteins. The real number of viral particles could be much higher in these samples and could be roughly estimated as the molecular weights of each viral protein are known and if the numbers of molecules per virus particle were documented for SARS-CoV-2. Here, we refer to the infectious dose as this is the most important parameter in terms of health concern, but the ratio of infectious particles in the nasopharyngeal swabs of patients may drastically differ from the purified virus fraction used here, and could even fluctuate during the course of the pathology. The strategy proposed for the analysis of these simili swabs consisted in a shotgun MS analysis based on a short acquisition of 20 min with a short LC gradient. For the clinical samples, we added an inclusion list of viral peptides in the MS method. The inclusion list allowed forcing the fragmentation of candidate viral peptide ions contained in the background matrix, even when they were not included in the top 20 from the data dependent acquisition method.

The shotgun strategy resulted in the detection of viral peptides in six out of the seven conditions tested for the simili swab experiment. From the five peptides of interest, GFYAQGSR and ADETQALPQR proved to be the most detectable and most sensitive in this background matrix, allowing proteotyping the virus up to the condition of 6 ng of viral material (7 PFU). One of the most interesting result was the omnipresence of peptide ADETQALPQR and its two miss-cleaved versions KADETQALPQR and KKADETQALPQR. These peptides were consistently detected in 30 out of 36 identifications in the six most abundant conditions from the simili swab experiment. Peptide KADETQALPQR was identified in all simili swabs from **Figure 2**. These results clearly show that ADETQALPQR, despite being prone to missed-cleavages, is one the most abundant and ionisable peptides and should be the main target for proteotyping SARS-CoV-2. This result was confirmed from the analysis of the clinical swab samples, since peptides ADETQALPQR, KADETQALPQR, and KKADETQALPQR were undoubtedly the most abundant in samples from COVID-19 patients (**Figure 4** and **Table 1**). In our previous work, we showed that this peptide sequence is also specific to SARS-CoV-2, but presented several variants among the available SARS-CoV-2 genomes (23). Therefore, when targeting this peptide for viral detection with MS/MS we can also take into account both its missed-cleaved versions, and its different variants.

Surprisingly, the high intensity peptide EITVATSR was only identified in simili swabs with high concentration of viral proteic material (**Figure 2**). By analyzing the MS and MS/MS spectra from these samples, we confirmed that this peptide co-eluted with another intense precursor from the background matrix that was fragmented simultaneously. The low MASCOT ion score attributed to these spectra hindered the confident identification of this peptide. This co-elution effect is most likely due to the use of the short chromatographic gradient, and one way to tackle it would be to use smaller isolation windows for fragmentation. This parameter was tested for the analysis of clinical swabs, but little or no improvement was observed. No MS/MS spectra were validated at FDR 1% for this peptide in swabs T7 and T8, even with the presence of a MS1 peak corresponding to this peptide in swab T8. This peptide is therefore problematic in this type of matrix and probably not suited for tracking SARS-CoV-2 in nasal swab samples with our specific experimental setup.

The distribution of the peptides along with the chromatogram from **Figure 3** shows that the two most detectable peptides GFYAQGSR and ADETQALPQR eluted in a narrow window of retention time between 15 – 16 min in simili swab samples. For the clinical swab samples, we observed that the retention time for these two peptides was 15.42±1.09 min for peptide ADETQALPQR, and 15.93±1.06 minutes for peptide GFYAQGSR as established with Skyline (**Table S7**). In the light of these new results, we argue that targeting peptides ADETQALPQR and GFYAQGSR with an extra short LC gradient of 3 min coupled to the enrichment of these hydrophilic peptides prior the LC injection could be one way to develop quick and robust assays for detection of the virus in clinical samples and gain in signal/noise ratio. Besides their high intensity, these peptides provide the needed specificity for a confident assay: peptide GFYAEGSR is highly conserved among different SARS-CoV-2 genomes, and peptide ADETQALPQR is specific to SARS-CoV-2. The simultaneous detection of these two peptides could provide therefore unequivocal evidence for the presence of the virus. Interestingly, a recent not yet published study showed the high potential of the same two peptides by using a targeted proteomics assay (19). The authors report limits of detection in the mid-attomole range corresponding to theoretically 10,000 SARS-CoV-2 particles in their specific experimental set-up.

Besides shortening the LC gradient to less than three min, sample preparation can also be optimized to develop more rapid peptidome preparation assays and remove too hydrophilic and too hydrophobic peptides that could saturate the chromatography column. Here, we performed a SDS-PAGE gel and in-gel proteolysis with trypsin to denature proteins and to remove any mass spectrometry-chromatography deleterious compounds that could be present in the nasal swab. This procedure is known to not be optimal as only 10% of the peptide material deposited on the gel is recovered. The literature is becoming rich in alternative sample preparation protocols for MS-based proteomics. For example, we recently proposed a proteotyping assay based on SP3 magnetic beads for protein purification and digestion in roughly 30 min (31). Being easily adapted to 96-well plates and robotization, SP3-based digestion is the method of choice for quick, high-throughput, and highly reproducible proteome digestions, as recently demonstrated (20, 32). Such sample preparation may further significantly increase the sensitivity of the tandem mass spectrometry proteotyping proposed in the present work. Furthermore, more sensitive instrument and MS acquisition modes could be tested to gain further sensitivity.

In conclusion, we tested in this study the potential of LC-MS/MS based methods for proteotyping SARS-CoV-2 in nasopharyngeal swabs. With a 20 min MS-acquisition window, we were able to identify and quantify several virus-specific peptides that allowed proteotyping the virus in simulated swabs and clinical swabs from COVID-19 patients. We argue that peptides ADETQALPQR (and its variant forms) and GFYAQGSR from the nucleocapsid protein are of utmost interest to develop quick and robust targeted assays for proteotyping the virus in nasopharyngeal swab samples. Further research must be done to validate their usefulness and their limits of detection in clinical samples, and develop the shortest possible pipeline.

## Author’s contributions

JA conceived the study with help from DG, GM, LG, and OP. GM and JCG performed the mass spectrometry experimental work. JPL and AS contributed the medical COVID-19 swab samples. FG, SD, and LB contributed the SARS-CoV-2 biological material. DG, LG, GM, OP and JA analysed the data. DG and JA wrote the manuscript with help from LG.

## Acknowledgements

The authors are indebted to Dr Silvia Meschi (National Institute for Infectious Diseases “Lazzaro Spallanzani” IRCCS, via Portuense 292, 00149 Rome, Italia) for making the Human 2019-nCoV strain 2019-nCoV/Italy-INMI1 (008N-03893) available. This publication was supported by the European Virus Archive goes Global (EVAg) project that has received funding from the European Union’s Horizon 2020 research and innovation programme under grant agreement N°653316. The authors are also grateful to the French Alternative Energies and Atomic Energy Commission (CEA), and the ANR program “Phylopeptidomics” (ANR-17-CE18-0023-01) that supported part of this study.

## Conflict of interest statement

The authors have declared no conflict of interest.

## Supplementary data (available upon request)

**Table S1.** List of *m/z* values of the inclusion list; **Table S2.** UNIPEPT taxonomical analysis of Swab R1, Swab R2, and simili swab 7R1-1pfu samples; **Table S3.** List of peptide-to-spectrum matches (PSMs) assigned from the simili SARS-CoV-2 swabs (FDR<1%); **Table S4.** List of proteins from the simili SARS-CoV-2 swabs and their spectral counts; **Table S5.** List of peptide-to-spectrum matches (PSMs) assigned in the clinical samples (FDR<1%); **Table S6.** List of proteins identified in the clinical samples and their spectral counts; **Table S7.** Skyline report with experimental data on precursor ions from the nine swabs from COVID-19 patients.

## Notes

https://www.ebi.ac.uk/pride/archive/projects/PXD019686

### Competing Interest Statement

The authors have declared no competing interest.

